# SCREAM: Single-cell Clustering using Representation Autoencoder of Multiomics

**DOI:** 10.1101/2025.10.03.680290

**Authors:** Panagiotis Chrysinas, Shriramprasad Venkatesan, Priya Ghanshyambhai Patel, Rudiyanto Gunawan

## Abstract

**Motivation:** Single-cell multiomics technologies offer unprecedented opportunities to study cellular heterogeneity. But, integrating information across different omics modalities remains a major challenge due to high dimensionality, sparsity, and modality-specific noise characteristics. To address this, we develop SCREAM (Single-cell Clustering using Representation Autoencoder of Multiomics), a novel deep learning framework for the robust integration and clustering of multi-modal single-cell data. SCREAM leverages Stacked Autoencoders (SAEs) to generate robust latent representations for each omics modality as well as for their fusion. Subsequently, borrowing Deep Embedding Clustering (DEC), SCREAM iteratively fine tunes the integrated mulitomics latent space and single-cell cluster assignments.

**Results:** We evaluated SCREAM against eleven state-of-the-art methods using SNARE-seq and CITE-seq datasets. In this benchmarking, SCREAM consistently demonstrated superior performance, yielding the highest or near-highest Adjusted Rand Index (ARI) and Normalized Mutual Information (NMI) scores on both datasets. These findings validate SCREAM as a highly accurate and robust approach for identifying cell types from multiomics data. Furthermore, its multiomics embeddings provides biologically meaningful latent representations for diverse downstream analyses.

**Availability:** SCREAM is available at http://www.github.com/cabsel/scream.

## INTRODUCTION

Recent advancements in single-cell sequencing technologies have revolutionized our ability to study cellular heterogeneity, an important aspect in understanding complex biological processes such as cancer progression, immune responses, and cellular differentiation (1–3). Scalable single-cell sequencing methods, such as 10X Chromium and Smart-seq2 (4), have paved the way for the development of technologies capable of profiling a wide range of biomolecules, collectively referred to as multiomics. These include single-cell chromatin accessibility sequencing (scATAC-seq) (5), DNA methylation (6), proteomics (7), and metabolomics (8). Recent innovations further enable the simultaneous measurement of multiple omics modalities within individual cells. Notable examples include sciCAR-seq (9), scCAT-seq (10), and SNARE-seq (11), which jointly profile chromatin accessibility (epigenomics) and mRNA expression (transcriptomics).

These single-cell multiomics technologies provide a more comprehensive view of cellular heterogeneity and the regulatory mechanisms underlying transcriptional processes compared to single-omics approaches (12–15). Transcriptomics captures gene expression by quantifying mRNA abundance in individual cells, while epigenomics measures DNA methylation states and chromatin accessibility, which are often associated with active transcription (16). Integrating transcriptomics and epigenomics data has been shown to inform the regulatory mechanisms driving cellular heterogeneity (17). Numerous tools have been developed for integrating bulk omics data (18), with a few of these, such as MOFA (19) and IntNMF (20), having been adapted for single-cell datasets. However, single-cell data differ fundamentally from bulk sequencing data (21) and present unique challenges, including high dimensionality, sparsity, and noise, as well as imbalances in dimensionality and sparsity across different omics. These challenges necessitate the development of methods specifically tailored for single-cell multiomics data analysis.

A number of clustering methods for single-cell multiomics data employ graph-based or statistical metrics to establish cell-to-cell similarity. Notably, Seurat’s Weighted Nearest Neighbor (WNN) strategy (22) constructs separate nearest neighbor graphs for each modality, then computes a weighted combination of these graphs based on each modality’s contribution to local cell similarity. This modality-weighted graph is used for clustering using standard graph-based algorithms (e.g., Louvain or Leiden). Similarly, CiteFuse constructs omics-specific similarity matrices using statistical metrics such as correlations or proportionality, which are then merged to give a fused multimodal cell-to-cell similarity matrix for clustering using standard algorithms (23). Several other methods such as SABEC (24), CALISTA (25), DIMMSC (26), and BREMSC (27) employ probabilistic models to evaluate cell likelihood based on their single or multi-omics profiles, followed by likelihood maximization to group similar cells under the assumption that cells in a cluster share the same model parameters.

Another major class of clustering methods proceeds by projecting different omics modalities into a shared low-dimensional latent space. One group of methods, including MOFA (19), its extension MOFA+ (28), and scAI (29), use factor analysis to construct shared latent factors across omics modalities, with clustering subsequently performed on the resulting factor matrix using standard techniques (*e.g*., *k*-means). Other methods, such as Liger (30) and Harmony (31), apply matrix factorization and alignment strategies to construct shared embeddings. All of the above methods adopt linear approaches that may be insufficient to capture the complex nonlinear relationships across omics modalities.

A family of recent nonlinear methods leverage machine learning strategies, specifically autoencoders, to construct latent space representations of single-cell multiomics data. Two related methods, including scCTClust (32) and MoClust (33), train separate modality-specific deterministic autoencoders, and apply divergence-based clustering strategy to the weighted sum of their latent vectors. Several methods are based on variational autoencoders (VAEs), including TotalVI (34), MultiVI (35), Cobolt (36), scMVAE (37), and Deep Cross-omics Cycle Attention (DCCA) (38), scMM (39). TotalVI and MultiVI use modality-specific encoders to generate joint embeddings, followed by Leiden clustering. However, separating the clustering step from dimensionality reduction may result in latent spaces that are suboptimal for clustering. The method scMVAE addresses this by coupling representation learning with cluster structure via a probabilistic Gaussian mixture in the latent space, encouraging cluster-aware embeddings. Cobolt employs a hierarchical multimodal VAE that integrates jointly profiled and modality-specific datasets into a single shared latent, explicitly modeling cross-modality correspondence and batch effects within the generative framework. In contrast, DCCA aligns omics-specific latent spaces using an attention-transfer mechanism, which can be powerful but often requires careful tuning and implicitly assumes near one-to-one cluster alignment. Lastly, scMM builds separate VAEs for different omics modalities, and then integrates their outputs using a mixture-of-experts approach to learn a unified latent representation for each cell.

In this study, we propose a latent space projection-based method, called *Single-cell Clustering using Representation Autoencoder of Multiomics* (SCREAM). SCREAM, illustrated in **Figure 1a**, consists of two main steps. First, the method independently trains two omics-specific Stacked Autoencoders (SAEs) (40) to learn low-dimensional latent representations for each omics modality. These modality-specific embeddings are then concatenated and fed into the Deep Embedding Clustering (DEC) algorithm (41), which performs representation-coupled clustering iteratively in the concatenated latent space. While DEC has been applied to single-cell clustering, its use has previously been limited to only scRNA-seq (42). We evaluate SCREAM on two SNARE-seq datasets, each comprising matched scRNA-seq and scATAC-seq data from the same cells, and on a CITE-seq dataset, profiling transcriptomes alongside cell-surface protein abundances. Our results show that SCREAM outperforms current single-cell multi-omics clustering methods, achieving more accurate and robust cell-type delineation than state-of-the-art baseline methods.

**Figure 1.**
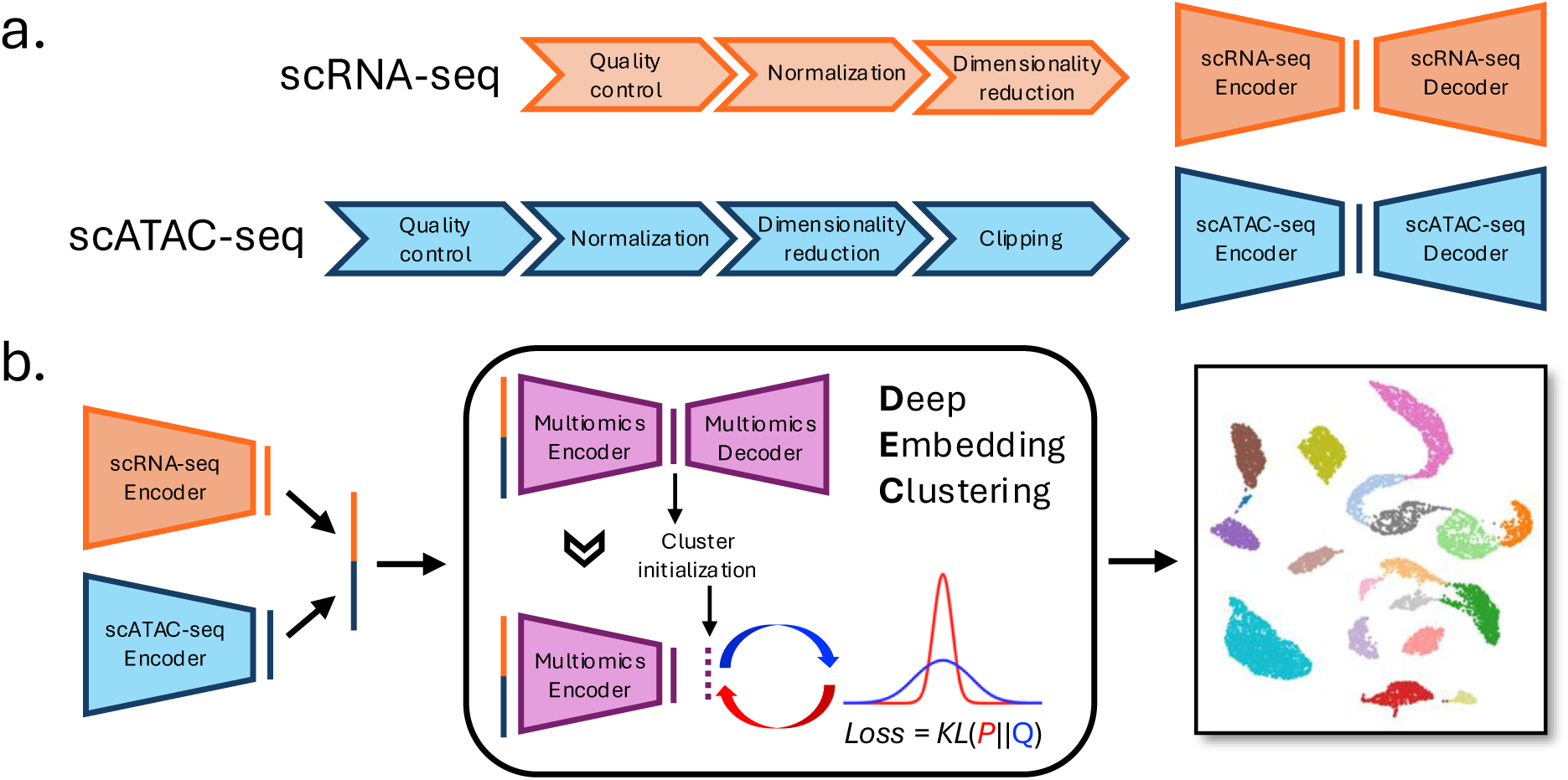
Overview of the SCREAM workflow. (a) Construction of stacked autoencoders (SAEs) for generating latent representations of individual omics modalities. Each modality is preprocessed according to established best practices. For scRNA-seq, preprocessing includes quality control, normalization, and dimensionality reduction using SCTransform (SCT) in Seurat. For scATAC-seq, preprocessing involves quality control, normalization using term frequency-inverse document frequency (TF-IDF), and dimensionality reduction via singular value decomposition (SVD) and clipping, also implemented in Seurat. The preprocessed data are then used as input for the training of respective SAEs. (b) Clustering in SCREAM. The encoders from (a) are used to generate latent representations for each omics modality. These representations are concatenated and subjected to single-cell multiomics clustering using the Deep Embedding Clustering (DEC) algorithm. DEC operates in two steps: first, a denoising SAE is trained for the concatenated multiomics representations; second, clustering is performed by iteratively optimizing the encoder parameters and cluster centroids through the construction of an auxiliary target distribution and the minimization of the Kullback–Leibler (KL) divergence between the soft cluster assignments and this target distribution

## METHODS AND MATERIALS

### Data Preprocessing

SCREAM is designed to integrate multiple single-cell omics datasets derived from the same cell population (i.e., single-cell multiomics). While its performance was demonstrated using scRNA-seq and scATAC-seq multiomics data from SNARE-seq, the approach can be adapted to other omics modalities. Preprocessing for each omics data type follows standard best practices, as illustrated in **Figure 1a** for scRNA-seq and scATAC-seq data.

#### Preprocessing of scRNA-seq Data

For scRNA-seq UMI count data, we performed normalization and variance stabilization using the sctransform (SCT) method implemented in Seurat (22). Following normalization, we filtered the dataset to retain only highly variable genes, with a default threshold of 3,000 genes.

#### Preprocessing of scATAC-seq Data

For scATAC-seq peak matrix data, we applied a modified version of the term frequency-inverse document frequency (TF-IDF) algorithm tailored for scATAC-seq data (43,44). The TF-IDF computation is based on two components:

1. Term Frequency (TF):

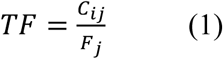

where for each cell in cell *j*, *C*_*ij*_ is the total number of counts for peak *i* and *F_j_* is the total number of counts.
2. Inverse Document Frequency (IDF):

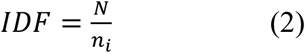

where *N* is the total number of cells, and *n_i_* is the total number of counts for peak *i* across all cells.

The TF-IDF matrix was then calculated as:

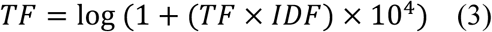

Next, we applied dimensionality reduction using the RunSVD routine from the Signac package (44), which employs partial Singular Value Decomposition (SVD) to convert peaks into Latent Semantic Indices (LSIs) (45). By default, RunSVD generates 50 LSIs, but we found that SCREAM achieved optimal performance when using 300 LSIs. Consistent with standard practice, we removed the first LSI component, as it typically captures technical variation in sequencing depth rather than biological variation (44). To further mitigate the impact of outliers, we clipped the LSI values to a range between −10 and 10.

### Stacked Autoencoder Training

SCREAM utilizes Stacked Autoencoders (SAEs) to generate latent representations for both individual and integrated omics data. An SAE consists of an encoder and a decoder, each comprising multiple hidden layers. **Figure 2** provides an example of an SAE with two hidden layers. The training process for an SAE involves two steps: pre-training and fine-tuning.

**Figure 2.**
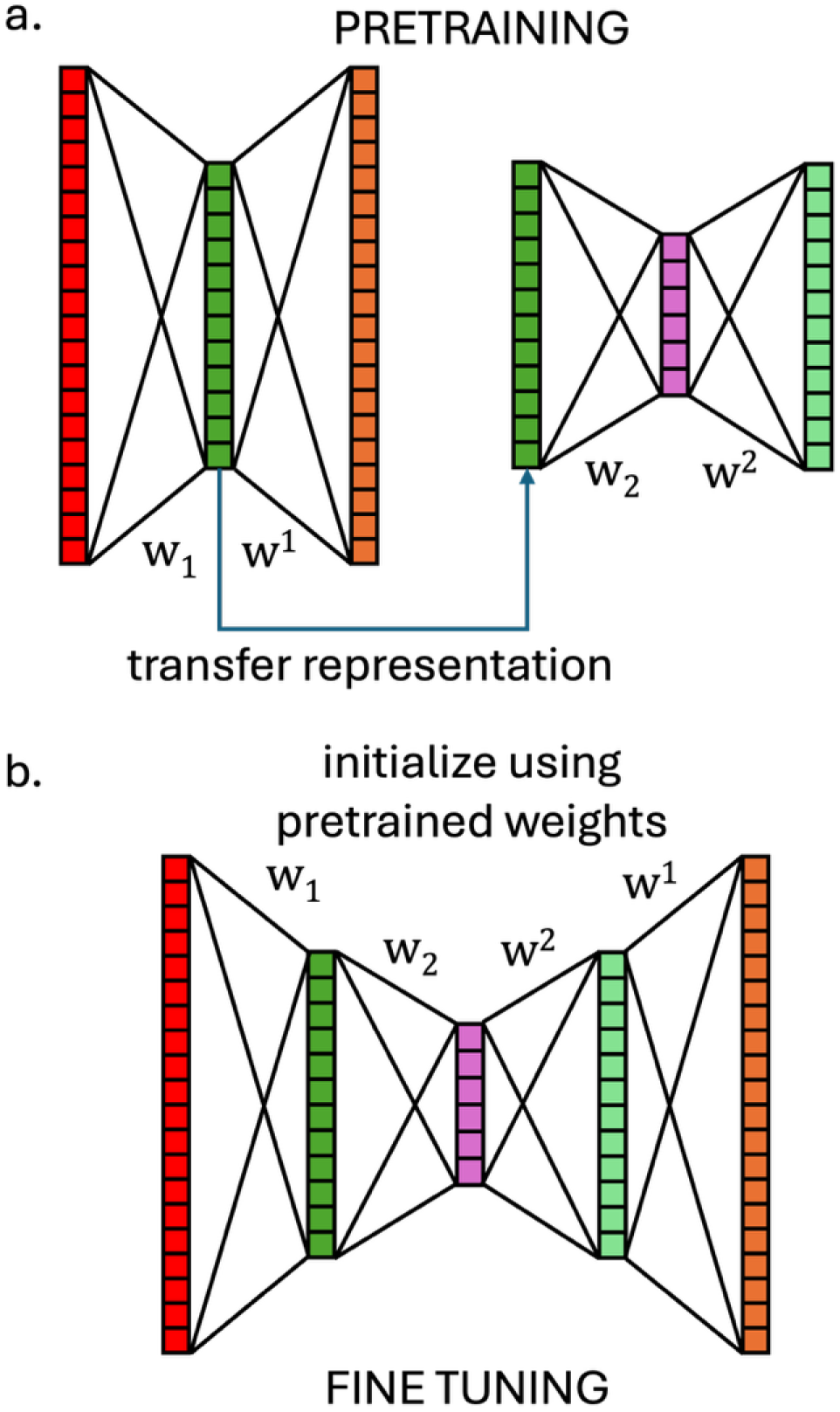
Training of a Stacked Autoencoder (SAE) with two hidden layers. (a) *Pre-training step*: ach layer of the SAE is trained sequentially as a denoising autoencoder. Random noise is injected into the input data via dropout, and the model is trained to reconstruct the original, noise-free input. The first layer encodes the input data into a compressed representation. (b) *Fine-tuning step*: The pre-trained encoder and decoder layers are stacked to form the full SAE, which is then fine-tuned end-to-end. During this phase, the model minimizes the reconstruction loss between the original input and the final output, allowing all layers to adjust their parameters simultaneously.

The pre-training step employs a sequential approach to train each autoencoder layer individually, starting from the outermost layer and progressing inward (see **Figure 2a**). During this stage, each autoencoder is trained as a denoising autoencoder: random noise is injected into the input data via dropout, and the model is trained to reconstruct the original, noise-free input. This denoising training encourages each layer to learn robust and meaningful features that are resilient to noise and irrelevant variation (40). The layer-wise training ensures that each layer learns useful representations independently and provides well-initialized parameters for the subsequent deep autoencoder.

After pre-training, the encoders and decoders from the individually trained autoencoders are assembled in reverse order of their layer-wise training to construct the full SAE (see **Figure 2b**). The entire SAE is then fine-tuned in an end-to-end manner by minimizing the reconstruction loss between the original input (without noise) and the final output. During this phase, all layers are updated simultaneously, allowing the model to adjust its parameters simultaneously to learn complex hierarchical patterns in the data. This end-to-end fine-tuning enhances the multilayer deep autoencoder to generate more accurate and meaningful latent representations.

### SCREAM Workflow

In the following, we describe the SCREAM workflow for integrating scRNA-seq and scATAC-seq multiomics datasets, noting that the framework is adaptable to other omics combinations following the same procedure. Let *X*_*RNA*_ ∈ ℝ^*n*×*p*^ and *X*_*ATAC*_ ∈ ℝ^*n*×*q*^ be the processed scRNA-seq and scATAC-seq data, respectively. Here, *n* denotes the number of cells in the dataset, while *p* and *q* correspond to the number of highly variable genes and latent semantic indices (LSIs), respectively (see *Data Preprocessing*).

The SCREAM framework consists of two main steps: (i) representation learning for each omics modality and (ii) deep embedding clustering (DEC) on the concatenated latent representations. In the first step, denoising stacked autoencoders (SAEs) are trained separately for each omics modality, and their encoders are used to generate latent embeddings. These embeddings, denoted *Z*_*RNA*_ ∈ ℝ^*n*×*p̃*^ and *Z*_*ATAC*_ ∈ ℝ^*n*×*q̃*^, are concatenated to form the fused representation:

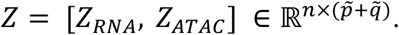

This concatenated embedding serves as input to the DEC algorithm (41), a method previously applied to single-cell transcriptomics (42) and here adapted for multiomics data integration and clustering. DEC proceeds in two phases:

1. Autoencoder pretraining: An additional autoencoder is trained on the fused embedding *Z*, following the denoising SAE framework. The encoder from this model yields the integrated multiomics latent representation, which is further fine-tuned during clustering.
2. Iterative clustering: Cluster centroids are initialized using Leiden clustering (46), but other methods such as *k*-means or Louvain can also be employed. DEC alternates between:

a. *Soft assignment*: Computing a similarity metric *q_ij_* between the embedding of the *i*-th cell *z*_*i*_ and the *j-*th cluster centroids *μ_j_* using the Student’s t-distribution as a kernel (47):

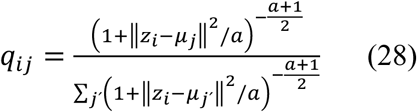

where *a* is the degrees of freedom of the Student’s t-distribution (*a* = 1 to avoid unnecessary learning).
b. *Clustering refinement*: Defining an auxiliary target distribution *P*:

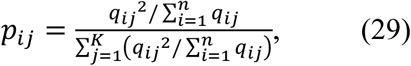

and minimizing the Kullback–Leibler (KL) divergence between *P* and *Q*:

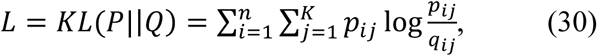

jointly optimizing the encoder weights and cluster centroids via stochastic gradient descent with momentum. Training continues until cluster assignment stability is reached or the maximum number of epochs is exceeded.

Several hyperparameters can influence SCREAM performance. In the representation learning stage, the number of hidden layers and latent dimensions for each SAE can be tuned to control denoising and compression within each omics modality. In the clustering stage, similar hyperparameters govern the fused autoencoder. Additionally, the clustering resolution used during initialization is critical, as it guides the refinement process and affects the granularity of final clusters. For hyper-parameter tuning, we used the Bayesian optimization tuner from the `keras-tuner` Python package. We employed the minimization of reconstruction loss by the autoencoders as the tuning objectives and performed 50 trials to identify the best set of hyper-parameters.

### Application of SCREAM to SNARE-seq datasets

SNARE-seq is a single-cell multimodal profiling technology that is capable of providing measurements of a cell’s transcriptome and its accessible chromatin. By integrating information about gene expression and chromatin accessibility, SNARE-seq provides insights into the regulatory mechanisms governing gene expression and their relationship with chromatin accessibility (11). We applied SCREAM to two SNARE-seq datasets to evaluate its performance.

#### CellLine dataset

The first dataset, referred to as the CellLine dataset, comprises 1,047 cells and includes both scRNA-seq and scATAC-seq data (11). These cells were derived from mixtures of cultured human BJ, H1, K562, and GM12878 cell lines. The scRNA-seq and scATAC-seq data were previously preprocessed by Zuo et al. (38), and we used their preprocessed data as inputs to SCREAM to enable direct comparisons with other methods tested on the same dataset.

For the scRNA-seq data, 500 highly variable genes (HVGs) were selected using the `FindVariableFeatures` routine from Seurat, which employs the variance-stabilizing transformation (VST) for feature selection (48). This process involves fitting a local polynomial regression (loess) to the logarithmic relationship between variance and mean, standardizing feature values using the observed mean and expected variance, and computing feature variance after clipping to a maximum value (default: square root of the number of cells). Meanwhile, for the scATAC-seq data, peaks within the gene body of the 500 HVGs and up to 100 kbp upstream were selected. The preprocessed CellLine data were then used as inputs to SCREAM, allowing for direct comparisons with other methods previously evaluated on this dataset.

#### 10XPBMC dataset

The second dataset, referred to as the 10XPBMC dataset, consists of human peripheral blood mononuclear cells (PBMCs) provided by 10X Genomics (49). This dataset contains 11,909 individual cells, which have been previously characterized and assigned to 20 cell types based on their transcriptome profiles (48). For data preprocessing, we followed the pipeline illustrated in **Figure 1a**. Quality control was performed based on the Seurat Weighted Nearest Neighbors (WNN) vignette (48). The quality control metrics included the number of detected molecules for each omics modality and the mitochondrial gene percentage. Specifically, we retained cells with count depths between 1,000 and 25,000 for scRNA-seq and between 5,000 and 70,000 for scATAC-seq. Additionally, cells with a mitochondrial gene percentage of 20% or higher were excluded. After preprocessing, the scRNA-seq data were reduced to the 3,000 most variable genes, while the scATAC-seq data were reduced to 299 latent semantic indices (LSIs). Cell type assignment was described in Hao *et al.* (48) and cells with a cell type prediction score less than 0.5 were removed. This resulted in a final dataset of 9966 high-quality cells.

### Application of SCREAM to CITE-seq dataset

CITE-seq is a single-cell multimodal profiling technology that enables simultaneous measurement of a cell’s transcriptome and surface protein (epitope) abundance using oligonucleotide-labeled antibodies (50). By capturing both gene expression and protein marker information, CITE-seq provides a more comprehensive view of cellular identity and heterogeneity. To evaluate SCREAM’s performance on a multimodal single-cell dataset different from SNARE-seq, we applied the method to a CITE-seq PBMC dataset.

The CITE-seq dataset, available from 10X Genomics, consists of 7,865 peripheral blood mononuclear cells (PBMCs) from a healthy donor. The preprocessed dataset was previously used for benchmarking MoClust and other methods and are available from the BREM-SC repository (27). Each cell was profiled with scRNA-seq and matched measurements of 14 TotalSeq-B antibodies (CD3, CD4, CD8a, CD14, CD15, CD16, CD19, CD25, CD45RA, CD45RO, CD56, CD127, PD-1, and TIGIT). Data preprocessing was performed as described by Yuan et al. (33), reducing the scRNA-seq data to 500 highly variable genes. Antibody expression was normalized with the centered log-ratio (CLR) transformation. We applied SCREAM to cluster the cells based on the integrated transcriptomic and protein epitope modalities. The original cell type annotations provided by 10X Genomics were used as the reference for evaluating clustering accuracy.

### Accuracy Assessment

The accuracy of SCREAM clustering results were evaluated by using two metrics: the normalized mutual information (NMI) (51) and the adjusted Rand index (ARI) (52). Both metrics are widely used in assessing clustering accuracy (33,38,42). Both metrics range between 0 and 1, where a value of 1 indicates perfect match with the ground truth.

The ARI measures the similarity between two clustering assignments while adjusting for chance. The ARI is evaluated as follows:

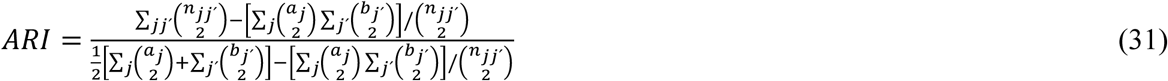

where *n*_*jj’*_, is the number of overlapping cells between cluster *j* in the reference (*i.e.,* cell type label) and cluster *j*’ in SCREAM’s clustering results. The variables *a_j_* and *b_j’_* denote the number of cells assigned to clusters *j* and *j*’, respectively. The ARI adjusts for the possibility of random clustering, making it a robust metric for evaluating clustering accuracy.

Meanwhile, the NMI quantifies the amount of information shared between the reference clustering and SCREAM’s clustering results. The NMI is evaluated as follows:

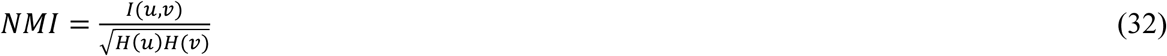

where *u* and *v* are the vectors of clustering assignment of the cells in the dataset (*i.e.*, cell type labels) and in SCREAM result, respectively. The term (*u*, *v*) is the mutual information of *u* and *v*, while (*u*) and *H*(*v*) are the entropies of *u* and *v,* respectively. NMI is a symmetric metric, meaning it treats the reference and predicted clustering equally, making it particularly useful for comparing clustering results with varying numbers of clusters.

## RESULTS

### SCREAM integrates and clusters single-cell multiomics data

SCREAM is a versatile framework designed for the integration and clustering of single-cell data, where multiple omics modalities (e.g., transcriptomics and epigenomics) are profiled from the same cells. The framework employs denoising Stacked Autoencoders (SAEs) in two key steps. First, denoising SAEs are trained independently for each omics modality to learn robust latent representations, effectively reducing noise and minimizing modality-specific biases. These latent representations are then concatenated to create a unified multiomics latent space. In the next step, the Deep Embedding Clustering (DEC) algorithm trains a denoising SAE on the concatenated latent representation, producing an integrated multiomics representation of the cells. The DEC process iteratively alternates between computing soft cluster assignments and minimizing the Kullback-Leibler (KL) divergence, progressively refining the multiomics latent space. This iterative optimization balances the contributions of different omics modalities, allowing SCREAM to capture shared and complementary information across modalities and generate biologically meaningful clusters.

In this study, we applied SCREAM to two publicly available SNARE-seq datasets: CellLine and 10XPBMC, and to a CITE-seq dataset (see **Methods and Materials**). To evaluate SCREAM’s clustering performance, we used two widely adopted metrics: the adjusted Rand index (ARI) and normalized mutual information (NMI).

### SCREAM achieves superior clustering performance on the CellLine dataset

Figure 3a gives the performance of SCREAM in clustering the CellLine dataset as a function of Leiden resolution, ranging from 0.1 to 1. The vertical dashed line marks the lowest Leiden resolution value (0.1) that produces the same number of clusters as the reference cell types (4 cell types). At this resolution, SCREAM achieved its highest ARI and NMI scores, demonstrating its ability to produce biologically accurate clusters.

**Figure 3.**
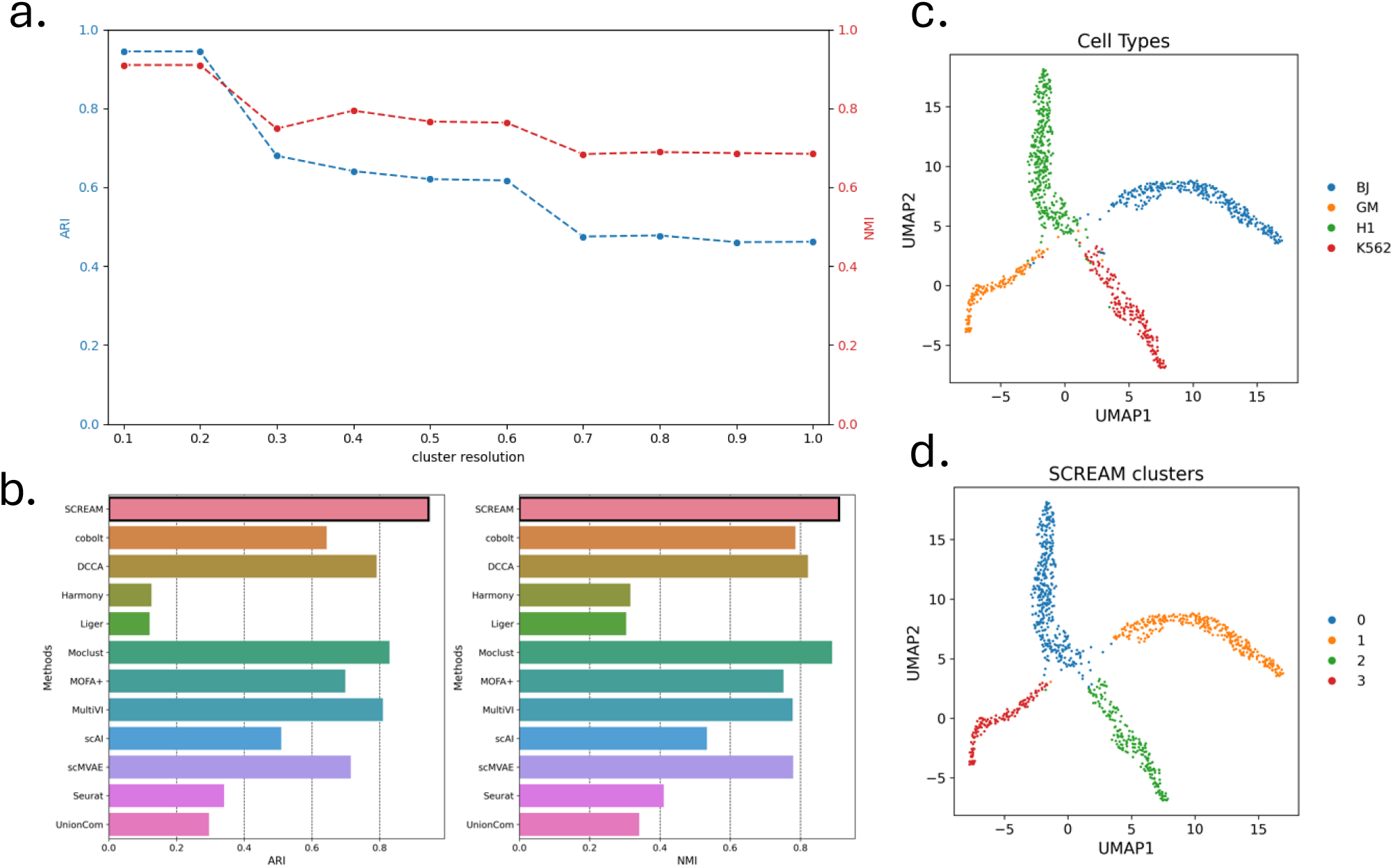
Comparative evaluation of SCREAM and existing methods on the CellLine dataset. (a) ARI and NMI scores for SCREAM across different Leiden resolution values. Bayesian optimization prescribed an RNA-seq autoencoder with hidden dimensions [1024, 256], an ATAC autoencoder with hidden dimensions [800, 12], and a fused autoencoder with hidden dimensions [1024, 256]. (b) Benchmark comparison of SCREAM with state-of-the-art clustering methods (also see **Supplemental Table S1**). All methods were implemented to give the same number of clusters as the reference cell types. (c, d) UMAP visualizations of SCREAM’s multiomics latent representation (Leiden resolution = 0.1), colored according to the reference cell types (c) and SCREAM’s clustering assignments (d).

Figure 3b compares SCREAM’s clustering performance with 11 other state-of-the-art clustering methods, including MoClust (33), Seurat V4 (48), MOFA+ (28), scAI (29), Cobolt (36), scMVAE (37), DCCA (38), MultiVI (35), Liger (30), and Harmony (31). The implementation details and results of these comparative methods were described in Yuan *et al*. (33). SCREAM’s performance was evaluated for the result from using a Leiden resolution of 0.1—the lowest resolution that generated the same number of cell type labels. SCREAM achieved the highest ARI score of 0.94 and NMI score of 0.91. Meanwhile, the second-best method was MoClust with an ARI score of 0.83 and a NMI score of 0.89. Among nonlinear methods, MultiVI gave the best performance, followed by scMVAE and cobolt. Linear methods generally performed worse than nonlinear strategies, with the exception of MOFA+ with similar accuracies as cobolt. Seurat, a widely used single-cell analysis toolbox, performed surprisingly poorly on this dataset. Methods employing alignment of multiomics latent space, including Liger, Harmony, and UnionCom, also performed poorly.

Figures 3c-d present the UMAP visualizations of SCREAM’s multiomics embeddings, showing a strong agreement between the reference cell types and SCREAM’s clustering results. For example, the cell types BJ, GM, H1 and K562 correspond almost perfectly to SCREAM clusters ID 1, 3, 0, and 2, respectively. This result highlights SCREAM’s ability to generate biologically meaningful clusters with high accuracy.

### SCREAM excels in clustering the 10XPBMC dataset

To further assess SCREAM’s performance, we applied it to the 10XPBMC SNARE-seq dataset that has significantly larger number of cells (9966 vs. 1,047 in CellLine) and cell types (19 vs. 4) than the CellLine dataset. Figure 4a shows the ARI and NMI scores as a function of Leiden resolution, showing a stable trend of accuracies across different Leiden resolutions. In the comparison below, we used SCREAM’s ARI and NMI from the clustering result using a Leiden resolution of 1.75, the lowest resolution that generated the same number of clusters as the reference.

**Figure 4.**
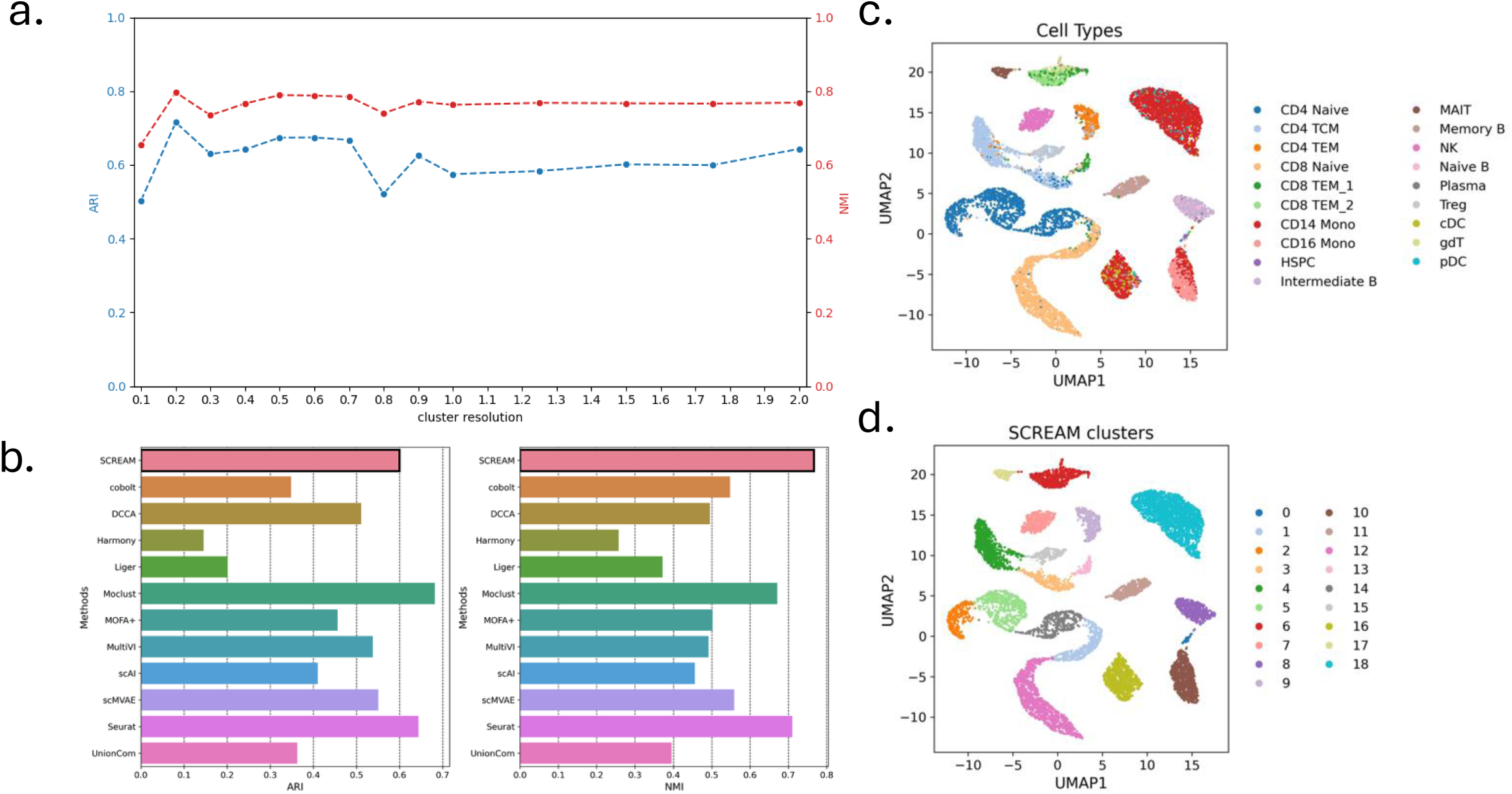
Comparative evaluation of SCREAM and existing methods on the 10XPBMC dataset. (a) ARI and NMI scores for SCREAM across different Leiden resolution values. Bayesian optimization prescribed RNA-seq, ATAC-seq, and fused autoencoders with the same hidden dimensions [1024, 256]. (b) Benchmark comparison of SCREAM with state-of-the-art clustering methods (also see **Supplemental Table S2**). All methods were implemented to give the same number of clusters as the reference cell types. (c, d) UMAP visualizations of SCREAM’s multiomics latent representation (Leiden resolution = 1.75), colored according to the reference cell types (c) and SCREAM’s clustering assignments (d).

Figure 4b compares SCREAM’s performance with the same state-of-the-art methods as in the above. SCREAM achieved the highest NMI score of 0.76 while achieving an ARI score of 0.59—a score that is better than the other methods except for MoClust (0.68) and Seurat (0.64). But we noted that SCREAM’s ARI and NMI peaked at a lower Leiden resolution of 0.2 that generated 7 clusters, reaching 0.72 and 0.80, respectively (see **Supplemental Figure S1**). As with the CellLine dataset, Figure 4b also shows nonlinear methods generally outperforming linear methods and latent alignment-based methods having the lowest accuracies.

Figures 4c-d present UMAP visualizations of SCREAM’s multiomics embeddings, illustrating a high concordance between the reference cell types and SCREAM’s clustering results. While the visual agreement is less striking than in the CellLine dataset, as anticipated with the larger number of cell types in the 10XPBMC dataset, distinct clusters corresponding to specific cell types are still apparent. For instance, NK cells, CD8 Naïve T cells, CD14 Monocytes, and memory B cells correspond to SCREAM clusters 12, 1, 18, and 11, respectively.

In summary, SCREAM demonstrated superior performance on the clustering of single-cell transcriptomic-epigenomic datasets, achieving the highest NMI score and the best or near-best ARI score among all methods. These results highlight SCREAM’s effectiveness and robustness in integrating and analyzing multiomics data, even in challenging scenarios with large datasets and numerous cell types.

### SCREAM outperforms in Clustering CITE-seq Dataset

To evaluate SCREAM’s performance on a different single-cell multiomics dataset, we applied it to the CITE-seq PBMC dataset (see Methods and Materials). Figure 5a shows the ARI and NMI scores across different Leiden resolutions. For consistency with the previous case studies, we selected SCREAM’s clustering result at the smallest Leiden resolution (0.1), which yielded the same number of clusters as the reference cell types.

**Figure 5.**
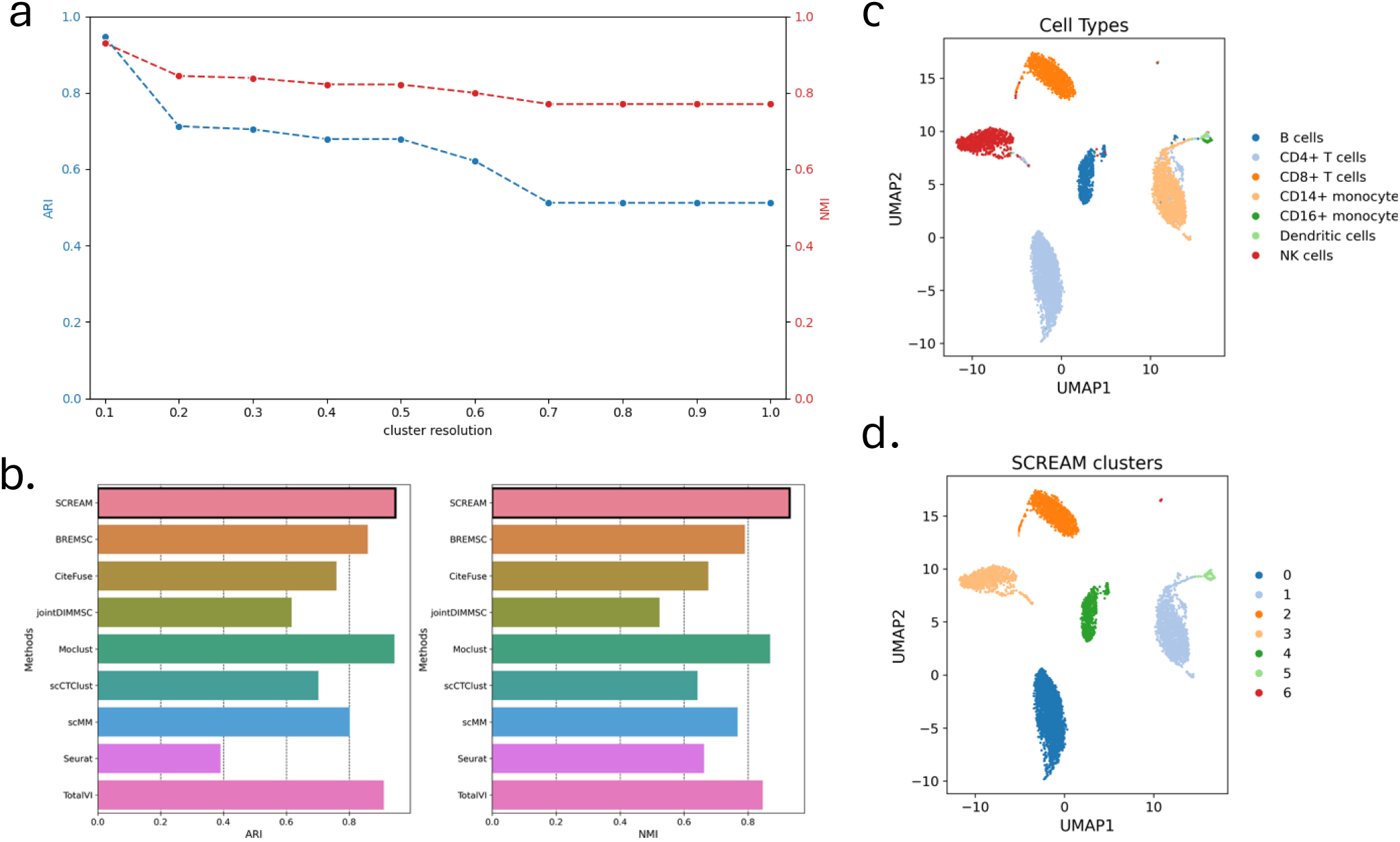
Comparative evaluation of SCREAM and existing methods on the CITE-seq PBMC dataset. (a) ARI and NMI scores for SCREAM across different Leiden resolution values. Bayesian optimization prescribed RNA-seq and ATAC-seq with hidden dimensions [50, 25], and fused autoencoder with hidden dimensions [1024, 256]. (b) Benchmark comparison of SCREAM with state-of-the-art clustering methods (also see **Supplemental Table S3**). All methods were implemented to give the same number of clusters as the reference cell types. (c, d) UMAP visualizations of SCREAM’s multiomics latent representation (Leiden resolution = 0.1), colored according to the reference cell types (c) and SCREAM’s clustering assignments (d).

Figure 5b benchmarks SCREAM against a panel of state-of-the-art clustering methods for CITE-seq data, including BREMSC (27), CiteFuse (23), DIMM-SC (26), MoClust (33), scCTClust (32), scMM (39), Seurat (22), and TotalVI (34). The implementation details and results of these comparative methods were described in Yuan *et al*. (33). SCREAM achieved the highest clustering accuracy, with an ARI of 0.95 and NMI of 0.93, outperforming all competing methods. MoClust ranked second, achieving ARI and NMI scores of 0.94 and 0.87, respectively. The VAE-based approaches TotalVI and scMM, along with the probabilistic framework BREM-SC, also performed relatively well. By contrast, Seurat and DIMM-SC gave the lowest accuracies on this dataset.

Figures 5c–d provide UMAP visualizations of SCREAM’s multiomics embeddings for the reference cell type and SCREAM’s clustering assignments, respectively. The latent representation learned by SCREAM clearly delineates major immune cell subpopulations with a strong correspondence between the reference cell type labels and inferred clusters. Specifically, CD4+ T cells, CD14+ monocytes, CD8+ T cells, natural killer cells, B cells, and dendritic cells, map onto SCREAM clusters 0 through 5, respectively. This close alignment demonstrates that SCREAM not only achieves superior quantitative accuracy but also recovers biologically meaningful clusters in CITE-seq data.

## DISCUSSION

This work addresses the challenge of integrating single-cell multiomics data to achieve a more comprehensive understanding of cellular functions. Single-cell multiomics technologies, which simultaneously measure multiple omics modalities from the same cell, offer unprecedented opportunities to uncover complex regulatory mechanisms. However, integrating these data remains challenging due to the high dimensionality, sparsity, noise, and imbalances in dimensionality, sparsity, and ultimately information encoded in different omics modalities. SCREAM was developed to address these challenges by employing a deep learning-based method, Deep Embedding Clustering (DEC), to fuse latent representations of multiple omics modalities and assign cells to clusters. SCREAM’s strength lies in its ability to balance and fuse multiomic information for clustering through the use of Stacked Autoencoders (SAEs), ensuring that both omics modalities contribute meaningfully to the final integrated representation. This balancing ensures that both omics contribute optimally to the clustering process, mitigating possible biases introduced by differences in dimensionality or sparsity.

The benchmarking results demonstrate SCREAM’s superior performance compared to other state-of-the-art methods on SNARE-seq and CITE-seq datasets. On the CellLine SNARE-seq and CITE-seq dataset, SCREAM achieved the highest ARI and NMI, outperforming the other state-of-the-art methods. Similarly, on the complex 10XPBMC dataset, SCREAM achieved the highest NMI and a second-highest ARI. These results highlight SCREAM’s robustness and accuracy in clustering single-cell multiomics data, even in challenging scenarios with large datasets and numerous cell types.

Like many deep learning-based methods, SCREAM’s performance depends on several hyperparameters. In the representation learning and clustering stages, we employed Bayesian strategy to optimize the SAE dimensions, performing 50 trials with the minimization of reconstruction loss as the tuning objective. Additionally, the Leiden clustering resolution used during initialization is critical, as this guides the refinement process and affects the number of final clusters. Since the optimal number of clusters is often not known a priori, we screened different

Leiden resolution values. Users can adopt similar approaches by systematically evaluating different resolution values, employing complementary techniques such as silhouette analysis, the elbow method, or gap statistics, or leveraging domain knowledge about expected cell types.

The broader implications of SCREAM may extend beyond clustering. Besides clustering, the multiomics latent representation generated by SCREAM (*i.e.*, the output of SAE encoder in DEC) may also be used for different downstream applications. This multiomics embeddings could, for example, be adapted for trajectory inference, enabling the reconstruction of dynamic cellular processes. When applied to SNARE-seq data, the fused representation could aid gene regulatory network reconstruction, capturing both transcriptomic and epigenomic signals to uncover regulatory relationships.

## CONCLUSION

In this work, we present SCREAM, a deep learning framework designed to address the challenges of integrating and clustering single-cell multiomics data. SCREAM follows a two-step approach: it first employs Stacked Autoencoders (SAEs) for latent representations of individual omics modality, and then applies a Deep Embedding Clustering (DEC) for multiomics fusion and clustering. Benchmarking on the CellLine and 10XPBMC SNARE-seq datasets, as well as the CITE-seq PBMC dataset, demonstrated that SCREAM consistently outperforms the state-of-the-art methods. The framework is flexible and broadly applicable to diverse single-cell multiomics datasets and analyses. Furthermore, its multiomics embeddings provide biologically meaningful latent representations that enable a wide range of downstream analyses.

## ACKNOWLEDGEMENT

The authors gratefully acknowledge funding support from the National Institute of Allergy and Infectious Diseases (grant number AI174080), the National Science Foundation (award number 1940162), and the National Heart, Lung, and Blood Institute (grant number HL103411).

