## Supplemental Figure S1 for "SCREAM: Single-cell Clustering using Representation Autoencoder of Multiomics"

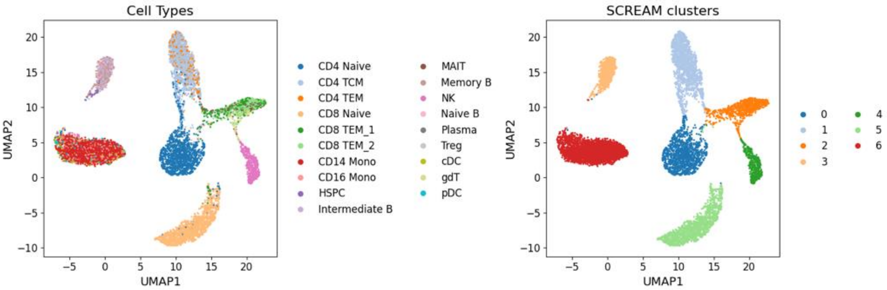


Supplemental Figure S1. UMAP of SCREAM multiomics embeddings for 10XPBMC dataset using Leiden resolution of 0.2.
